# Proposed Methodology for Reducing Bias in Structural MRI Analysis in the Presence of Lesions: Data from a Pediatric Traumatic Brain Injury Cohort

**DOI:** 10.1101/2023.02.12.528180

**Authors:** Daniel Griffiths-King, Adam Shephard, Jan Novak, Cathy Catroppa, Vicki A. Anderson, Amanda G. Wood

## Abstract

Traumatic brain injury can lead to multiple pathologic features, including brain lesions, which are visible on magnetic resonance imaging (MRI). These resulting heterogenous lesions can present a difficulty for several standard approaches to neuroimaging, resulting in bias and error in subsequent quantitative measurements. Thus, cases presenting with lesions on MRI may be excluded from analyses, biasing samples across the research field. We outline a potential solution to this issue in the case of Freesurfer, a popular neuroimaging tool for surface-based segmentation of brain tissue from structural MRI. The proposed solution involves two-steps, a) Pre-processing: Enantiomorphic Lesion-Filling and b) Post-processing: Lesion Labelling. We applied this methodology to 14 pediatric TBI cases which presented with lesions on T1w MRI. Following qualitative inspection of these cases after implementation of the approach, 8 out of 14 cases were retained as being of sufficient quality. In brief, we have presented here an adapted pipeline for processing structural MRI (sMRI) of patients who have experienced a TBI using the Freesurfer software package. This approach aims to mitigate potential lesion-induced biases that exist beyond the locality of the pathological tissue, even in the contralesioned hemisphere.

---

Structural MRI (sMRI) can be utilised to estimate functionally relevant brain ‘damage’ after a traumatic brain injury (TBI), primarily through the quantification of the morphometry of brain regions (see [1] for review). These approaches may be more sensitive to subtle effects of injury on the brain compared to routine visual reporting by neuroradiologists. Therefore, these methods may better allow the understanding of the neuroanatomical basis of later impairment.

The accuracy of these methods. however, are biased by errors introduced during the automated-processing of sMRI containing gross anatomical lesions and/or pathology. Essentially, morphometric measures generated for these cases may not be biologically valid for two main reasons; a) gross pathology (such as encephalomalacic regions) or pathological voxel intensities (such as gliosis or oedema) can either render boundaries undetectable or discontinuous [2-5], or b) systematic biases introduced by the presence of pathology on the Freesurfer pipeline (i.e. contralesional hemisphere biases [6] or atlas registration biases [3, 4]). These potential errors make it difficult to ascertain whether differences between control and patient morphology are due to an injury-related pathology or due to inaccuracies in morphometric measures specific to patients with gross lesions [1].

Historically, studies of paediatric TBI (pTBI) have excluded cases with major pathology present on MRI (for instance [7]) due to these potential processing biases. However, this limits the utility of previous research with the exclusion of these patients risking a systematic bias in sampling. Given that the location and extent of focal lesions to the brain following a pTBI are seemingly insufficient to fully explain post-injury neuropsychological deficits [8] (i.e. following early brain injury impaired executive function occurs irrespective of injury factors such as lesion location [9, 10]) Inclusion of these lesion cases in research may increase accuracy of prognostic quantitative models and ensure they generalise to the full spectrum of pathology [1, 2, 5, 6]. Therefore, approaches and/or methodologies that are robust to the presence of lesions are necessary for future studies.

In a recent paper, Diamond and colleagues [2, 5] identified and outlined a potential methodology with which to ‘optimise’ structural segmentation of sMRI for patients with TBI. This utilised the Freesurfer pipeline, an automated approach to the surface-based structural segmentation of T1w MRI. Diamond and colleagues’ [2, 5] approach involves the manual labelling of tissue where the reconstructed surfaces pass through cortical lesions. However, this post-processing approach, which results in very focal edits to the surface reconstruction, does not address global algorithmic biases indicated by the presence of lesions. For instance, in a recent study, we identified that the presence of simulated lesion pathology, resulted in a small but systematic bias in the contralesional hemisphere, and the magnitude of this bias is seemingly associated with voxel intensities within this pathology [6]. Diamond and colleagues’ approach [2, 5] will not account for this bias.

In the current paper, we highlight a potential adjustment to the Freesurfer pipeline to mitigate some of the observed-issues in surface-based parcellation of the cortex in the presence of traumatic lesions, particularly the bias in the surface-placement of the contralateral hemisphere to the lesion.

## Methods

The data used in the current experiment are a subset of an existing dataset of children who have experienced a TBI between the ages of five and 16 years of age. 114 patients with pTBI were recruited between 2007 and 2010 into a study on ‘Prevention and Treatment of Social Problems Following TBI in Children and Adolescents’. More detailed descriptions have been published elsewhere [11-13]. In brief, children with TBI were recruited on presentation to the Melbourne Royal Children’s Hospital’s emergency department. Patients were eligible if they: i) were aged between five and 16 years at the time of injury, ii) had recorded evidence of both a closed-head injury and also two post-concussive symptoms (such as headaches, dizziness, nausea, irritability, poor concentration), iii) had sufficient detail within medical records to determine injury severity (e.g., Glasgow Coma Scale (GCS; Teasdale and Jennett [14]), neurological and radiological findings), iv) had no prior history of neurological or neurodevelopmental disorder, non-accidental injuries or previous TBI, and v) were English speaking.

### MRI Acquisition

MRI were acquired sub-acutely after injury (<90 days post-injury). MRI were acquired at 3T on a Siemens Trio scanner (Siemens Medical Systems, Erlangen, Germany) using a 32-channel matrix head coil. The acquisition included a sagittal three-dimensional (3D) MPRAGE [TR = 1900 ms; TE = 2.15 ms; IR prep = 900 ms; parallel imaging factor (GRAPPA) 2; flip angle 9 degrees; BW 200 Hz/Px; 176 slices; resolution 1 × 1 × 1 mm] and sagittal 3D T2-FLAIR non-selective inversion preparation SPACE (Sampling Perfection with Application-optimised Contrast using different flip-angle Evolution) [TR = 6000 ms; TE = 405 ms; inversion time (TI) = 2100 ms; water excitation; GRAPPA Pat2; 176 slices; 1 × 1 × 1 mm resolution matched in alignment to the 3D T1w sequence].

### Lesion Delineation and production of lesion masks

A trained rater (JN) visually inspected participant’s MRI for pathology, scrolling through contiguous axial slices of the 3D T1w and FLAIR images independently. Identified lesions were segmented manually (by JN) by drawing binary lesion masks on each of the T1w and FLAIR MRI scans using the ROI editor tool in MRtrix3.0 [15]. A second rater (AS) visually confirmed these masks. For this study, only the lesion masks drawn on the T1w MRI scan were utilised/necessary.

TBI lesions are typically extremely heterogenous in appearance on MRI scans [16]. Increasingly, white matter hyperintensities (WMH i.e. Leukoaraiosis) and enlarged perivascular spaces (EPVS i.e. Virchow-Robin spaces) are recognised as potential biomarkers for an increased risk of later emerging diseases/diagnoses [16-19]. Consequently, we also segmented these abnormalities.

For lesion segmentation, the following criteria were applied; i) abnormality visible on >3 contiguous axial slices (i.e. ≥ 1.5 mm), ii) visible WMHs should appear hyperintense on FLAIR and hypointense on T1w MRI [19], iii) visible EPVS should appear hypointense on both T1w and FLAIR MRI, and be tubular shaped depending on lesion orientation [19]. Examples of these lesion masks can be seen in Figure 1A+B.

**Figure 1.**
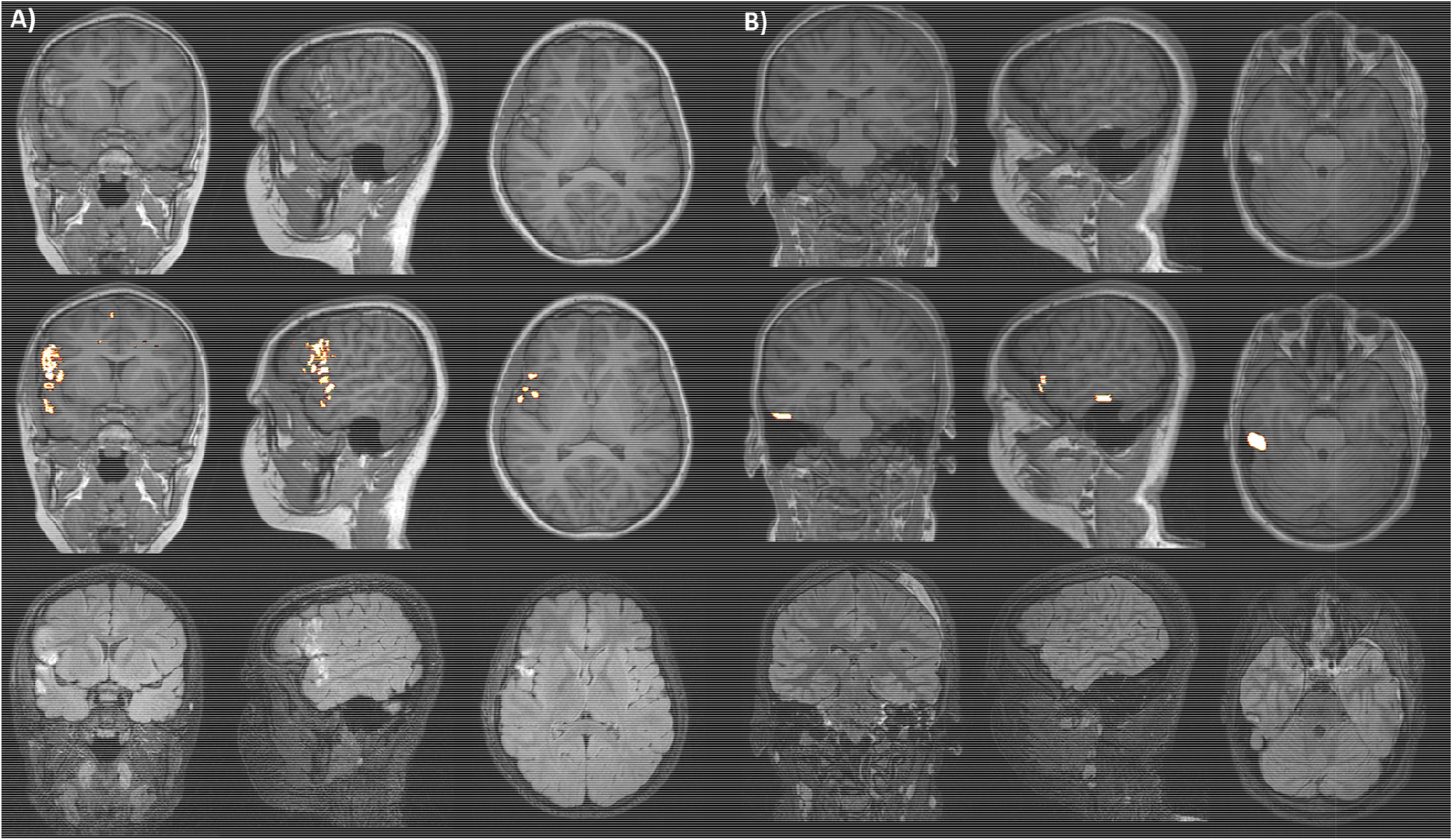
sMRI of two (A + B) patients with TBI with lesions. Top row: Unedited T1w image with visible gross pathology. Middle row: T1w image with overlaid binary lesion mask (mask is interpolated for visual purposes). Bottom row: Unedited FLAIR image.

### Proposed pipeline for processing sMRI with visible pathology

The current paper utilises a new approach to Freesurfer segmentation in the presence of focal lesions in the cortical GM ribbon, involving both pre- and post-processing procedures.

#### a) Pre-processing: Enantiomorphic Lesion-Filling

Lesion masks were used for pre-processing of MRI, to perform anatomically-informed lesion-filling, using the enantiomorphic approach of Nachev, Coulthard [20]. Briefly, this approach robustly registers the lesioned hemisphere to the contra-lesioned hemisphere and ‘fills’ the lesioned voxels (indicated by the lesion-mask) with subject-level, ‘healthy-appearing’ signal intensities from the homologous region in the contra-lesional hemisphere. The output is an MRI with approximately-typical T1w voxel-intensities, in place of the lesioned tissue. This step was conducted using the normalisation tool of the BCBlab (Brain Connectivity and Behaviour) [21]. We only performed these lesion-filling processes for those cases with frank GM lesions. Some recent evidence suggests that filling approaches for white matter lesions results in no changes to Freesurfer derived volume estimates [22]. This, and the fact that geometric inaccuracies due to WMH can be corrected using manual editing approaches as per Freesurfers’ guidelines, means that we focus on an approach to tackle GM lesion.

The enanteomorphically filled T1w image was then processed using the standard Freesurfer (6.0) cortical surface segmentation pipeline (using the –FLAIRpial commands) [23]. By processing this image rather than the original T1w MRI, we mitigate potential contrast-induced errors that may contribute to lesion-induced error/bias in structural segmentation, even in the contralesional hemisphere. An example of the resultant surfaces can be seen in Figure 2.

**Figure 2.**
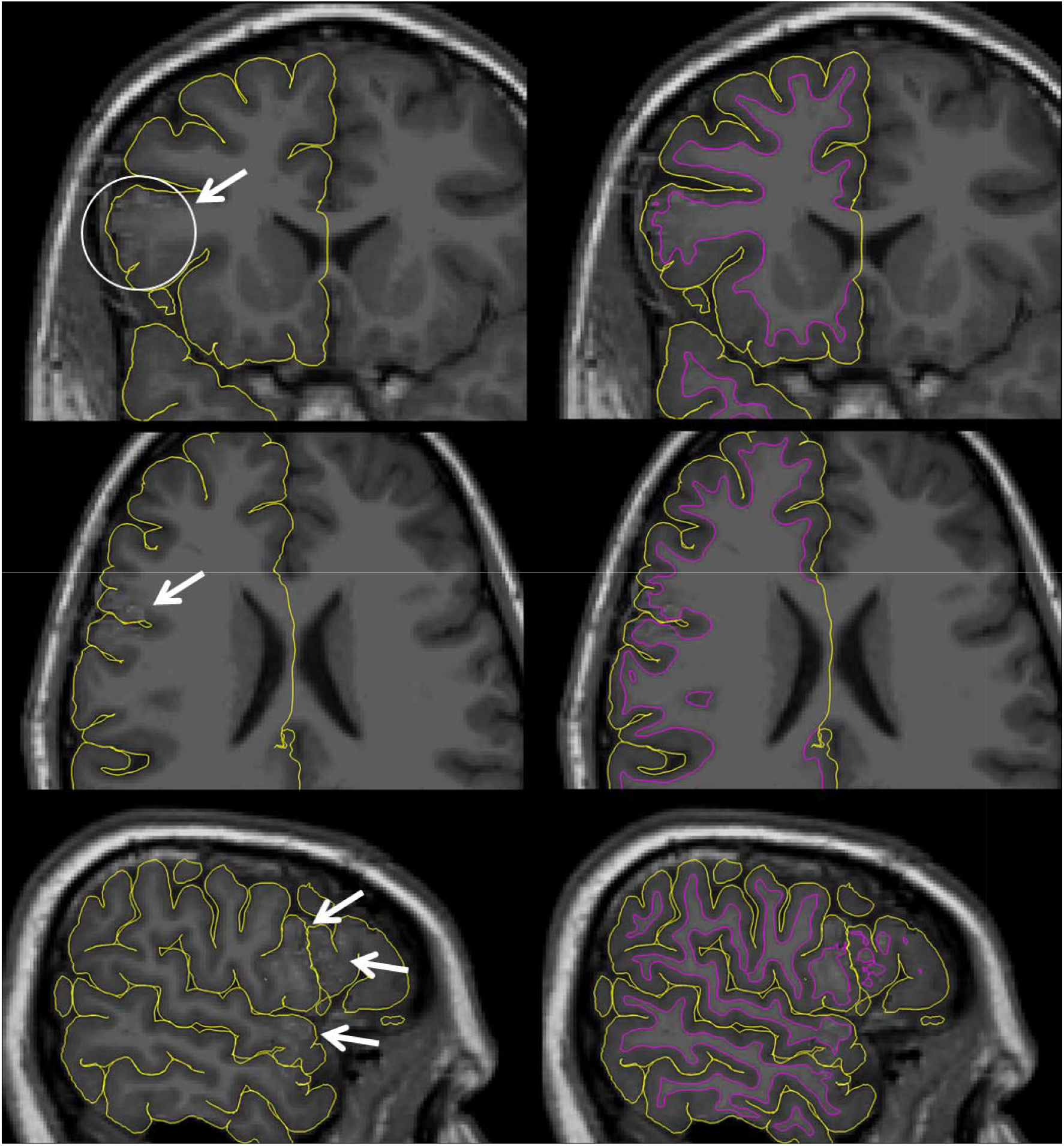
Freesurfer plotted surfaces overlaid on enanteomorphically filled T1w sMRI. Arrows highlight the filled areas. First column: Only pial surface visualised. Second column: Both pial and white surfaces visualised. The original lesion is identified in the MRI displayed in Figure 1A.

#### b) Post-processing: Lesion Labelling

Post-processing of the produced surface segmentations was also conducted. Lesion masks were projected onto the generated cortical surface vertices and the projected lesion ROI was filled (to avoid holes due to voxel-vertex mismatches). These approaches were adapted from scripts made available by the Multi-centre Epilepsy Lesion Detection (MELD) project [24, 25].

Individual-subject surface parcellations were masked using these surface projected lesion masks. Thus, region labels completely or partially occluded by lesion tissue were overwritten with the lesion label. Morphometric measures (such as cortical thickness, volume, etc) were calculated using standard Freesurfer approaches but, due to relabelling, no measures will be taken from tissue which is a) lesioned within the original image and b) filled with estimated voxel intensities in the enanteomorphically filled T1w images. For those regions that are completely occluded by the lesion label, morphological measures are reported as zero however, these can be recoded as ‘not a number’ (NaN) to ensure that they are not included in analyses and bias results.

The output of this pipeline is therefore cortical morphometric estimates for ROIs not contaminated by lesion tissue or the wider error associated with the processing of lesioned T1w images. A visual depiction of this can be seen in Figure 3.

**Figure 3.**
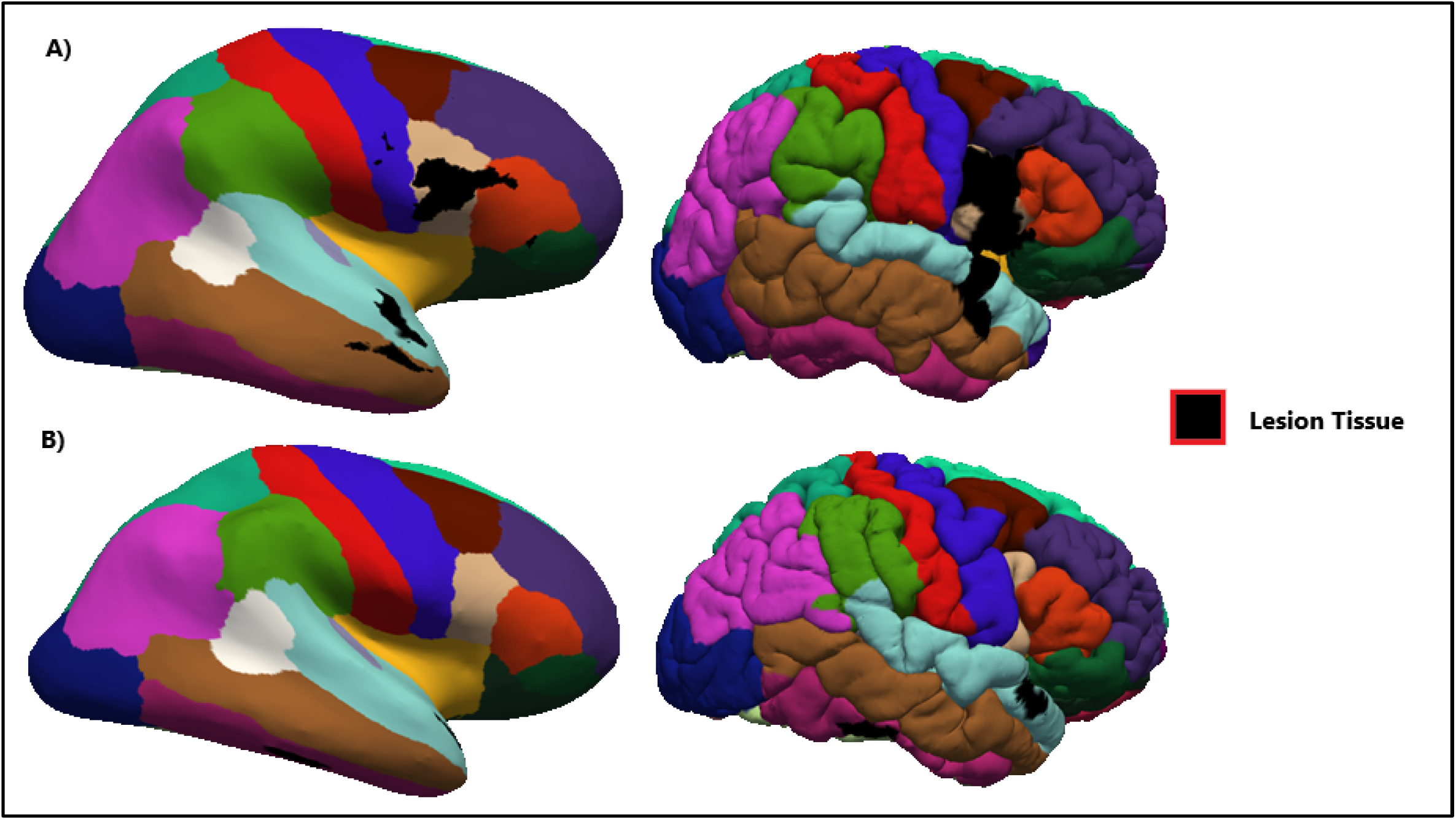
ROI parcellation based upon the Desikan-Killany atlas [26] for two lesion cases projected onto both the inflated (first column) and regular (second column) surface models for each subject. The lesion label can also be seen Subjects A) and B) relate to the corresponding subjects in Figure 1.

### Quality Assessments

Quality was visually assessed for all cases, based upon the delineation of both pial and white matter surfaces generated by Freesurfer. This allowed identification of cases where manual edits needed to be undertaken, and were carried out per standard Freesurfer protocols.

## Results

### Lesion Identification

Of the pTBI cases (N=114), we identified n=14 as having visible pathology definable as lesion tissue within the cortical GM ribbon. These cases were selected and underwent our lesion correction procedure

### QA

On initial visual inspection of the output of the lesion-correction procedure, two cases were initially excluded, due to the bilaterality of contusions in near-homologous regions, leading to unsuccessful lesion filling using the enanteomorphic approach. Five cases required no manual edits, although two of these had poor reconstruction focal in relation to the filled lesion tissue. However, as per Diamond and colleagues [2, 5], to reduce subjectivity in manual edits for these areas, no edits were undertaken, as these would later be labelled as lesion tissue. Seven cases underwent manual edits to improve surface placement. Of these, three cases were acceptable after editing. The four cases which were rejected were due to surface reconstruction issues that were related to motion present within the image, rather than due to the lesion correction procedure. The final number of cases processed with the lesion correction procedure was eight (out of 14).

### Post-Hoc Volume analyses

We conducted post-hoc analyses to assess whether the lesion correction methodology impacted cortical volumes and cortical thickness in the contralesioned hemisphere. Differences in both measures were found in both the contralesioned and lesioned hemispheres when comparing cases which have been corrected with the lesion pipeline versus those that have not been corrected with the lesion pipeline. Further details and figures can be found in supplementary materials. This suggests that that this method may in fact be correcting contralesioned hemisphere biases introduced by gross GM pathology found in [6].

## Discussion

The current paper highlights an alternative method to Diamond and colleagues [2, 5] for optimising the structural segmentation of sMRI in individuals with pTBI, specifically designed to mitigate the more global segmentation biases which are introduced in these lesioned cases [6]. This approach is being utilised in our lab enable the inclusion of lesioned cases into studies of the neuroanatomical correlates of cognitive impairment post paediatric TBI. The benefit of such methodologies is to simultaneously a) increase power to detect relevant case-control differences or brain-behaviour relationships and b) increase generalisability of findings to a wider spectrum of pathology. These benefits come about via the ability to include cases with visible pathology on MRI which previously would be precluded from analysis pipelines.

As outlined above, our approach differs from that of Diamond and colleagues [2, 5] in one major aspect. This is specifically the pre-processing of cases using an enanteomorphic filling approach. This was to tackle global biases in morphometric measures beyond the site of specific pathology (such as the contralesioned hemisphere) which may not be reduced by local correction methods. However, it may be argued that Diamond and colleagues’ [2, 5] approach better meets the first benefit outlined above, to increase statistical power through inclusion of cases. This is because, whilst our approach led to the inclusion of 8/14 cases which would have otherwise be excluded in future analyses, Diamond and colleagues [5] retained 87/98 MRIs for which Freesurfer surfaces were successfully generated and corrected using their methodology. However, this is not a direct comparison. It is important to remember that, in this study, cases were also excluded were removed for typical reasons not associated with the lesion, in this case motion artefact. This is unsurprising given the fact that this study utilised a paediatric population where movement artefact is more common [27], whereas Diamond [2, 5] investigated an adult cohort.

Whilst Freesurfer is primarily an automated tool for the processing of sMRI and generating surface-based models of cortical morphometry, these surface models can be and are frequently utilised further in the analysis pipeline of functional MRI and diffusion MRI studies of TBI. Therefore, effective methods to ensure inclusion of cases with visible lesions on MRI and reduce the biases that these lesions can introduce into these automated pipelines will have wide-reaching implications for the field. It is also important to note that, whilst TBI pathology is particularly heterogeneous, the effects of visible pathology on neuroimaging pipelines is not limited to TBI and thus the methods outlined here may also find use in other neurological disease (i.e. multiple sclerosis or tuberosclerosis).

This pipeline has been used in our lab in published works [28] allowing the inclusion of cases that typically may not have been able to be reliably included in research paradigms. A recent paper has outlined a similar approach termed “virtual brain grafting (VBG)” [29], which has subsequently been utilised in structural connectomic approaches to understanding the individualised effects of TBI on the brain [30]. Given the VGB approach was validated on “synthetic” MRI, it would be prudent to test the two methodologies head-to-head in future research.

## Limitations

One limitation of the current approach is its ability to handle bilateral lesions in homologous brain regions, put simply lesions to both hemispheres in approximately similar anatomical locations. This is because the enanteomorphic filling approach would attempt to ‘fill’ abnormal voxel intensities (the lesion) with abnormal, lesion voxel intensities from the ipsilateral hemisphere, rather than healthy tissue as intended. Two cases were excluded from the current study for this reason. Adopters of this approach must also be aware that, for the largest lesions, statistical analysis approaches, that robustly deal with missing data may be required. Fully occluded ROIs will be labelled as NaN, which is likely to be treated as missing data. to include them as zero makes an implicit assumption about the underlying tissue, that it is functionally and neuroanatomically irrelevant. These are important methodological considerations for the application of these approaches.

One potential issue with both proposed approaches is the reliance on accurate segmentations or ‘masks’ of lesion present on MRI. This can be difficult and time consuming, requires considerable expertise to be considered as ‘gold standard’ in the field (normally a trained neuroradiologist) with no ‘ground truth’ with which to truly assess performance. This can make approaches requiring such lesion masks prohibitive to; larger studies with a greater number of lesions to segment; labs without such neuroradiological expertise and most importantly clinical applications/practice where considerable ‘pre-processing’ would be required for new and incoming cases.

It is difficult to ascertain whether the approach outlined by Diamond and colleagues, or the approach outlined in the current study best optimises the Freesurfer pipeline for use in TBI cases with visible pathology, as there exists no ‘ground truth’ in these circumstances. Both approaches likely go some way in addressing the potentially biases introduces when Freesurfer is used to process MRI of patients with TBI and visible gross pathology. However, we would argue that our approaches goes further, trying to mitigate further biases which we have observed previously in these types of analyses [6]. Future work directly compare these methodologies using a more pragmatic approach, such as evaluating which method allows us to recover the most accurate predictions of cognitive functioning post-injury, or even predict injury-severity. These may be more clinically useful assessments of these methodologies in the absence of ‘ground truth’.

In brief, we have presented here an adapted pipeline for processing sMRI of patients who have experienced a TBI using the Freesurfer software package. This approach aims to mitigate potential lesion-induced biases that exist beyond the locality of the pathological tissue, even in the contralesioned hemisphere.

## Acknowledgments and Funding

The authors thank Dr Jan Novak and Dr Adam Shephard for their technical assistance in providing the lesion masks necessary for this study. This work was supported by a European Research Council (ERC) - Consolidator Grant (ERC-CoG) to A.G.W [grant number 682734]. This work was conducted whilst DGK was supported by a studentship from Aston University, School of Life and Health Science. and a Birmingham Childrens’ Hospital Research Foundation Grant (BCHRF) to Dr Sukhvir Wright and A.G.W. DGK is currently funded by a grant from Aston University, College of Health and Life Sciences to J.N.

## References

1. King, D.J., et al., A systematic review of cross-sectional differences and longitudinal changes to the morphometry of the brain following paediatric traumatic brain injury. Neuroimage Clin, 2019. 23: p. 101844.

2. Edlow, B., et al., Optimizing the Accuracy of Cortical Volumetric Analysis in Traumatic Brain Injury. Journal of Neurotrauma, 2019. 36(13): p. A62–A62.

3. Goh, S.Y., et al., Neuroinformatics challenges to the structural, connectomic, functional and electrophysiological multimodal imaging of human traumatic brain injury. Front Neuroinform, 2014. 8: p. 19.

4. Irimia, A., et al., Neuroimaging of structural pathology and connectomics in traumatic brain injury: Toward personalized outcome prediction. Neuroimage Clin, 2012. 1(1): p. 1–17.

5. Diamond, B.R., et al., Optimizing the accuracy of cortical volumetric analysis in traumatic brain injury. MethodsX, 2020. 7: p. 100994.

6. King, D.J., et al., Lesion Induced Error on Automated Measures of Brain Volume: Data From a Pediatric Traumatic Brain Injury Cohort. Front Neurosci, 2020. 14: p. 491478.

7. Serra-Grabulosa, J.M., et al., Cerebral correlates of declarative memory dysfunctions in early traumatic brain injury. Journal of Neurology Neurosurgery and Psychiatry, 2005. 76(1): p. 129–131.

8. Bigler, E., Quantitative magnetic resonance imaging in traumatic brain injury. The Journal of head trauma rehabilitation, 2001. 16(2): p. 117–134.

9. Anderson, V., et al., Children’s executive functions: are they poorer after very early brain insult. Neuropsychologia, 2010. 48(7): p. 2041–50.

10. Jacobs, R., A.S. Harvey, and V. Anderson, Are executive skills primarily mediated by the prefrontal cortex in childhood? Examination of focal brain lesions in childhood. Cortex, 2011. 47(7): p. 808–24.

11. Anderson, V., et al., Social competence at 6 months following childhood traumatic brain injury. J Int Neuropsychol Soc, 2013. 19(5): p. 539–50.

12. Anderson, V., et al., Social Competence at Two Years after Childhood Traumatic Brain Injury. J Neurotrauma, 2017. 34(14): p. 2261–2271.

13. Catroppa, C., et al., Social and Behavioral Outcomes following Childhood Traumatic Brain Injury: What Predicts Outcome at 12 Months Post-Insult? J Neurotrauma, 2017. 34(7): p. 1439–1447.

14. Teasdale, G. and B. Jennett, Assessment of Coma and Impaired Consciousness -Practical Scale. Lancet, 1974. 2(7872): p. 81–84.

15. Tournier, J.D., F. Calamante, and A. Connelly, MRtrix: Diffusion tractography in crossing fiber regions. International Journal of Imaging Systems and Technology, 2012. 22(1): p. 53–66.

16. Bigler, E.D., et al., Heterogeneity of brain lesions in pediatric traumatic brain injury. Neuropsychology, 2013. 27(4): p. 438–51.

17. Adams, H.H., et al., Rating method for dilated Virchow-Robin spaces on magnetic resonance imaging. Stroke, 2013. 44(6): p. 1732–5.

18. Dubost, F., et al., Enlarged perivascular spaces in brain MRI: Automated quantification in four regions. NeuroImage, 2019. 185: p. 534–544.

19. Wardlaw, J.M., M.C. Valdés Hernández, and S. Muñoz-Maniegax, What are white matter hyperintensities made of? Relevance to vascular cognitive impairment. Journal of the American Heart Association, 2015. 4(6): p. e001140.

20. Nachev, P., et al., Enantiomorphic normalization of focally lesioned brains. Neuroimage, 2008. 39(3): p. 1215–1226.

21. Foulon, C., et al., Advanced lesion symptom mapping analyses and implementation as BCBtoolkit. Gigascience, 2018. 7(3): p. 1–17.

22. Guo, C., et al., Repeatability and reproducibility of FreeSurfer, FSL-SIENAX and SPM brain volumetric measurements and the effect of lesion filling in multiple sclerosis. European Radiology, 2019. 29(3): p. 1355–1364.

23. Fischl, B., FreeSurfer. Neuroimage, 2012. 62(2): p. 774–81.

24. Adler, S., et al., MELD Protocol 4 - Lesion Masking. 2018: protocols.io. doi:10.17504/protocols.io.n9udh6w.

25. Adler, S., et al., Novel surface features for automated detection of focal cortical dysplasias in paediatric epilepsy. Neuroimage Clin, 2017. 14: p. 18–27.

26. Desikan, R.S., et al., An automated labeling system for subdividing the human cerebral cortex on MRI scans into gyral based regions of interest. Neuroimage, 2006. 31(3): p. 968–80.

27. Pardoe, H.R. R. Kucharsky Hiess, and R. Kuzniecky, Motion and morphometry in clinical and nonclinical populations. Neuroimage, 2016. 135: p. 177–85.

28. King, D.J., et al., Structural-covariance networks identify topology-based cortical-thickness changes in children with persistent executive function impairments after traumatic brain injury. NeuroImage, 2021. 244.

29. Radwan, A.M., et al., Virtual brain grafting: Enabling whole brain parcellation in the presence of large lesions. NeuroImage, 2021. 229: p. 117731.

30. Imms, P., et al., Exploring personalized structural connectomics for moderate to severe traumatic brain injury. Network Neuroscience, 2022: p. 1–24.

