## Supplemental Materials for "Proposed Methodology for Reducing Bias in Structural MRI Analysis in the Presence of Lesions: Data from a Pediatric Traumatic Brain Injury Cohort"

**Supplementary Materials**

**MRI Acquisition**

MRI were acquired as part of an existing research protocol described elsewhere (Anderson et al., 2013; Anderson et al., 2017; Catroppa et al., 2017). The MRI acquisition sequence specifically consisted of a sagittal three-dimensional (3D) MPRAGE [TR] = 1900 ms; TE = 2.15 ms; IR prep = 900 ms; parallel imaging factor (GRAPPA) 2; flip angle 9 degrees; BW 200 Hz/Px; 176 slices; resolution 1 × .5 × .5 mm], sagittal 3D T2-w non-selective inversion preparation SPACE (Sampling Perfection with Application-optimised Contrast using different flip-angle Evolution) [TR = 6000 ms; TE = 405 ms; inversion time (TI) = 2100 ms; water excitation; GRAPPA Pat2; 176 slices; 1 × .5 × .5 mm resolution matched in alignment to the 3D T1-weighted sequence].

**Post-Hoc Volume analyses**

To assess the proposed methodology we compared contralesional hemisphere volumes in seven of the eight processed cases. One case was excluded for having bilateral lesions which did not affect the enanteomorphic filling process (these bilateral lesions were found in non-homologous regions), but did not allow for the comparison of lesioned versus contralesioned hemispheres.

We compared hemispheric volumes of the eight cases processed with the standard Freesurfer pipeline and processed with the proposed lesion correction pipeline. In order to remove the bias of manual editing between the two sets of cases [2], we compared automated volumes where manual editing had not occurred. We compared both contralesioned and lesioned hemisphere volumes to assess whether the enanteomorphic filling process was leading to bilateral changes to the morphometric measures. Contalesioned hemisphere was selected based upon visual inspection of the lesion mask to assess the hemisphere contralateral to the major GM pathology. If the lesion correction procedure were successfully correcting for the expected bias in the contralesioned hemisphere, we would expect to see changes to measures within this hemisphere. Due to small sample sizes in this cohort, we did not perform quantitative statistics on these measures however visualisations can be seen in Supplementary Figure 1.

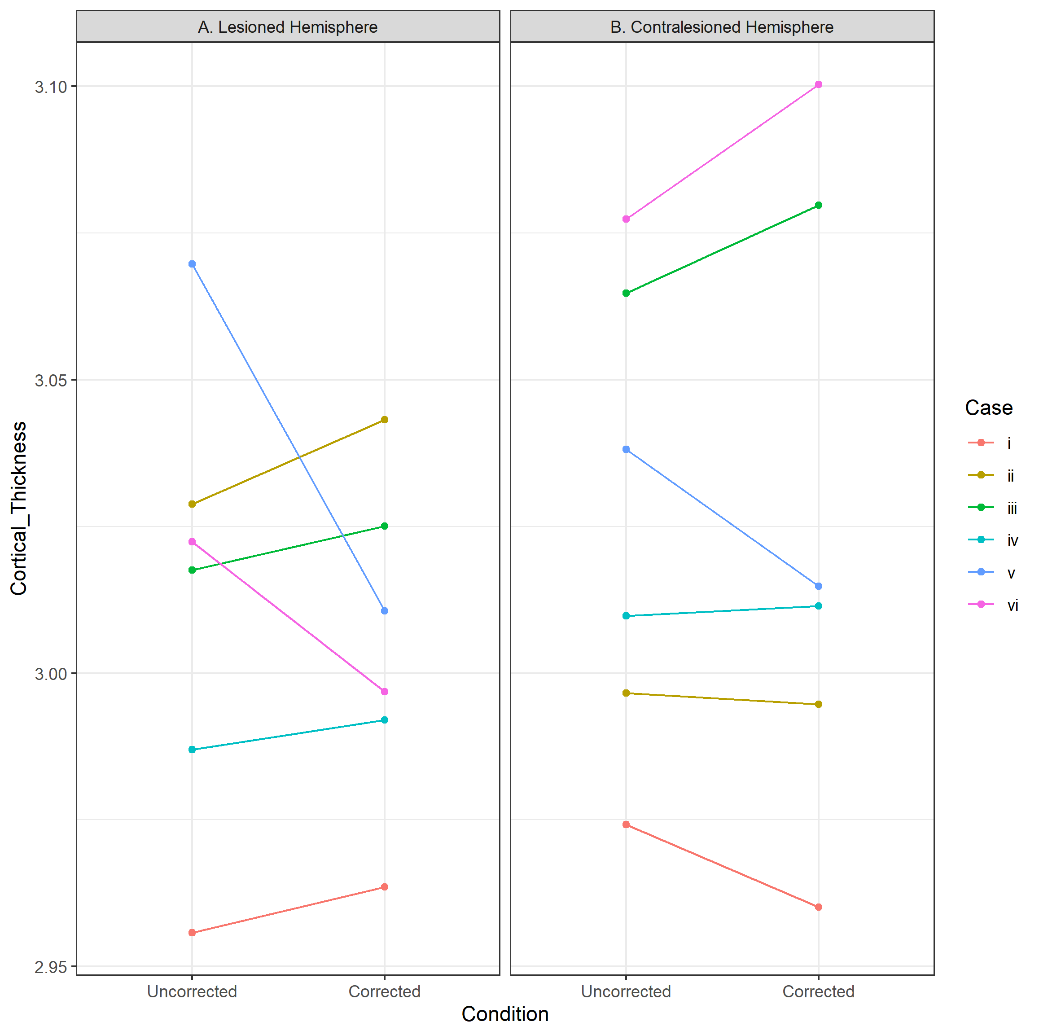

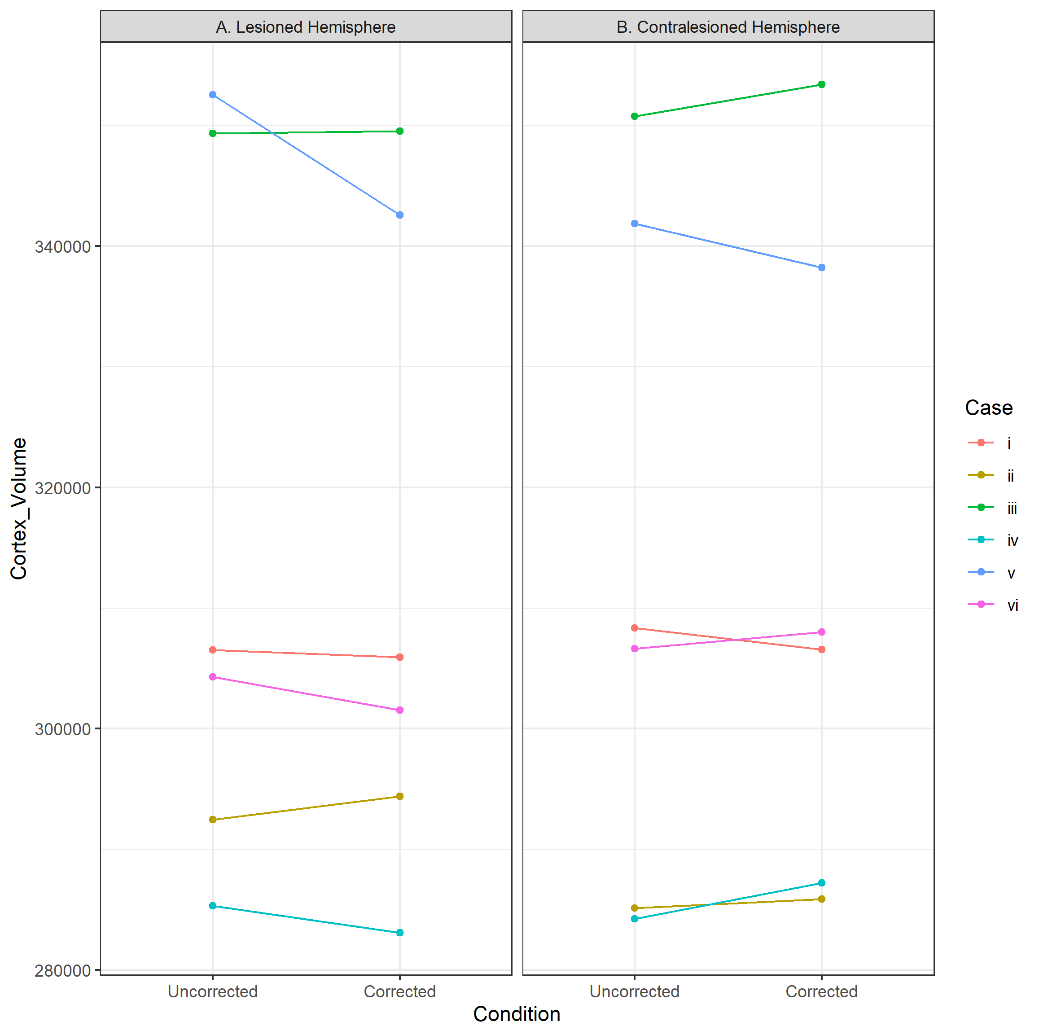

Supplementary Figure 1. Visualisations of the difference in i) Cortical Thickness and ii) Cortex Volume between those MRI processed with our pipeline (corrected) and those that have not (uncorrected) for both the lesioned and contralesioned hemispheres. This suggests that the lesion filling process seemingly has an effect on the contralesioned hemisphere, although this effect does not seem uniform in magnitude or direction, likely due to differing lesion characteristics [6]
